# Functional Brain Imaging and Targeted Lesion Studies Using Manganese-Enhanced MRI and Focused Ultrasound in Non-Conventional Model Species

**DOI:** 10.1101/2025.06.05.658064

**Authors:** Pierre Estienne, Gwenaël Pagé, Benoit Larrat, Kei Yamamoto, Luisa Ciobanu

## Abstract

Linking behavior to its neuroanatomical basis in non-conventional model species remains a significant challenge due to the scarcity of imaging and molecular tools. Commonly used approaches such as electrophysiological recordings rely on precise stereotaxic atlases or species-specific antibodies, while optogenetics requires transgenic lines which are largely unavailable beyond classical model organisms (e.g., mice, rats, zebrafish). Moreover, surgical lesion studies, a staple for verifying brain structure and behavior relationships, are logistically complex in species lacking atlases or living in aquatic environments. Here, we present a protocol integrating Manganese-Enhanced Magnetic Resonance Imaging (MEMRI) and MR-guided High-Intensity Focused Ultrasound (HIFU) to overcome these limitations, which we demonstrate in the convict cichlid (*Amatitlania nigrofasciata*), a teleost fish lacking conventional neuroscience tools. MEMRI enables non-invasive, sub-millimeter resolution mapping of brain activity during behavior, and HIFU facilitates precise, surgery-free lesioning of targeted regions, adaptable to species without stereotaxic atlases. This combined approach offers a versatile, broadly applicable framework for linking brain structure and behavior in non-model organisms, advancing evolutionary and comparative neuroscience.

## INTRODUCTION

Correlating behavior with its neuroanatomical substrate is challenging in non-conventional model species, for which few imaging or molecular tools are available, significantly impeding progress in neuroethology and comparative neuroanatomy^1^. A classical method for linking a measurable behavior to its neural basis involves electrophysiological recordings of brain activity to identify the regions engaged during the behavior. However, this approach is possible only when one already has a good candidate brain region to target the electrodes in a precise manner. In addition, electrophysiology is dependent on extremely precise stereotaxic brain atlases, which are available only in a limited number of model species such as macaque monkeys^2^, rats^3^, mice^4^, pigeons^5^ or zebra finches^6^.

As a post-mortem verification, immediate early genes (IEG), revealed either at their transcription stage through in-situ hybridization or at their translation stage through immunohistochemistry, are typically used to assess brain activity after a behavioral task in non-model species^7–11^. However, finding the right antibody can be difficult, as most antibodies are manufactured to work in model species and may show limited efficacy in other species^12,13^. An additional challenge when studying brain activation through immediate early genes is knowing where to look: whole brain IEG can be a labor-intensive process in large-brained species as it requires the sectioning, mounting, microscopic imaging and counting of neurons on a large scale. IEG also requires euthanizing the animal to sample the brain, which might not be possible in species where large numbers of individuals are unavailable.

More recently, in frequently used model species such as mice, rats, or zebrafish, imaging and molecular tools have become available, with transgenic lines allowing for calcium imaging, optogenetics and chemogenetic ablations^14–18^. Nonetheless, these are not available for less commonly used animals (birds, reptiles, other fish, or invertebrates such as cephalopods). This technical limitation significantly impedes progress in neuroethology and comparative neuroanatomy, as most studies use a very limited range of model species as a result^19–21^.

Beyond detecting brain activity, lesion studies to assess impairments in task performance have been crucial to the field of comparative neuroanatomy to verify the role of specific brain structures in behavior^22–25^. Given the lack of available techniques, surgical lesions are thus the preferred option for non-model species. However, the vast majority of species still do not have a stereotaxic brain atlas. Moreover, surgical lesions can be particularly difficult to perform when targeting deep structures, requiring going through non-targeted structures to reach the region of interest, at the risk of damaging those structures as well. Finally, basic surgical technique can be difficult to adapt to aquatic species, where working in a sterile environment to prevent infections is very difficult.

The lack of resources to measure brain activity and assess the impact of lesions on behavior in non-model species limits the breadth of biological questions that can be answered, at a time when the benefits of using non-conventional models in a comparative, evolutionary approach have become clear^21,26^.

Magnetic resonance imaging (MRI) has seen limited use in the fields of neuroethology and comparative neuroanatomy, in part due to the relatively high cost of preclinical MR scanners. However, as the cost of preclinical MR scanners decreases, MRI shows potential for non-invasive functional brain imaging in a wide range of species. Manganese-Enhanced MRI (MEMRI) has been shown to work in both vertebrates^27,28^ and invertebrates^29^ as a reliable reporter of neuronal activity. Mn^2+^ ions enter neurons via Ca_v_ 1.2 channels preferably, and accumulate in projection terminals^30^. In addition to MEMRI, lesion studies can be conducted without the need for surgery using MR-guided High-Intensity Focused Ultrasound (HIFU) by focusing non-ionizing ultrasonic waves at precisely targeted brain regions^31^. However, even though the technique is well documented to perform thermal ablations in rodent brains^32^, it requires adjustments to the constraints of non-model species.

Here, we describe a protocol combining MEMRI and HIFU in the convict cichlid (*Amatitlania nigrofasciata*), a teleost fish for which most traditional neuroscience tools are not available. We further describe how to adapt both techniques to any species, the only limitation being the available imaging space in the MRI scanner and the size of the ultrasound transducer available to the researcher. MEMRI allows for the identification of brain regions activated during a behavior of interest in a non-invasive manner with sub-millimeter resolution, and HIFU allows to perform reliable lesion studies in the absence of stereotaxic brain atlases, or in species for which clean and effective surgical lesions are not possible (e.g., aquatic species).

Together, these two techniques could find broad utility in the fields of neuroethology and comparative neuroanatomy, enabling more comprehensive studies of non-model animal species.

## RESULTS

### Manganese-Enhanced Magnetic Resonance Imaging (MEMRI)

In order to visualize neuronal activity in the brain of the convict cichlid (*Amatitlania nigrofasciata*), manganese-enhanced magnetic resonance imaging (MEMRI) was used^29^. MEMRI uses manganese, a magnetic resonance contrast agent, to label active neurons. Manganese was administered intraperitoneally (Fig. 1a) (n=22), as we determined that intramuscular injections, at least in convict cichlids, did not allow for a rapid diffusion of manganese to the brain. A dose of 50 mg/kg of MnCl_2_, well tolerated in fish, and providing good MR contrast was used (Fig. 2d). Previous studies in mice^33,34^ and our own unpublished data also show that this dosage appears well tolerated in endotherms as well, including mammals (i.e, rats) and birds (cockatiels, unpublished data), making it an ideal starting point for future studies using other species.

**Figure 1.**
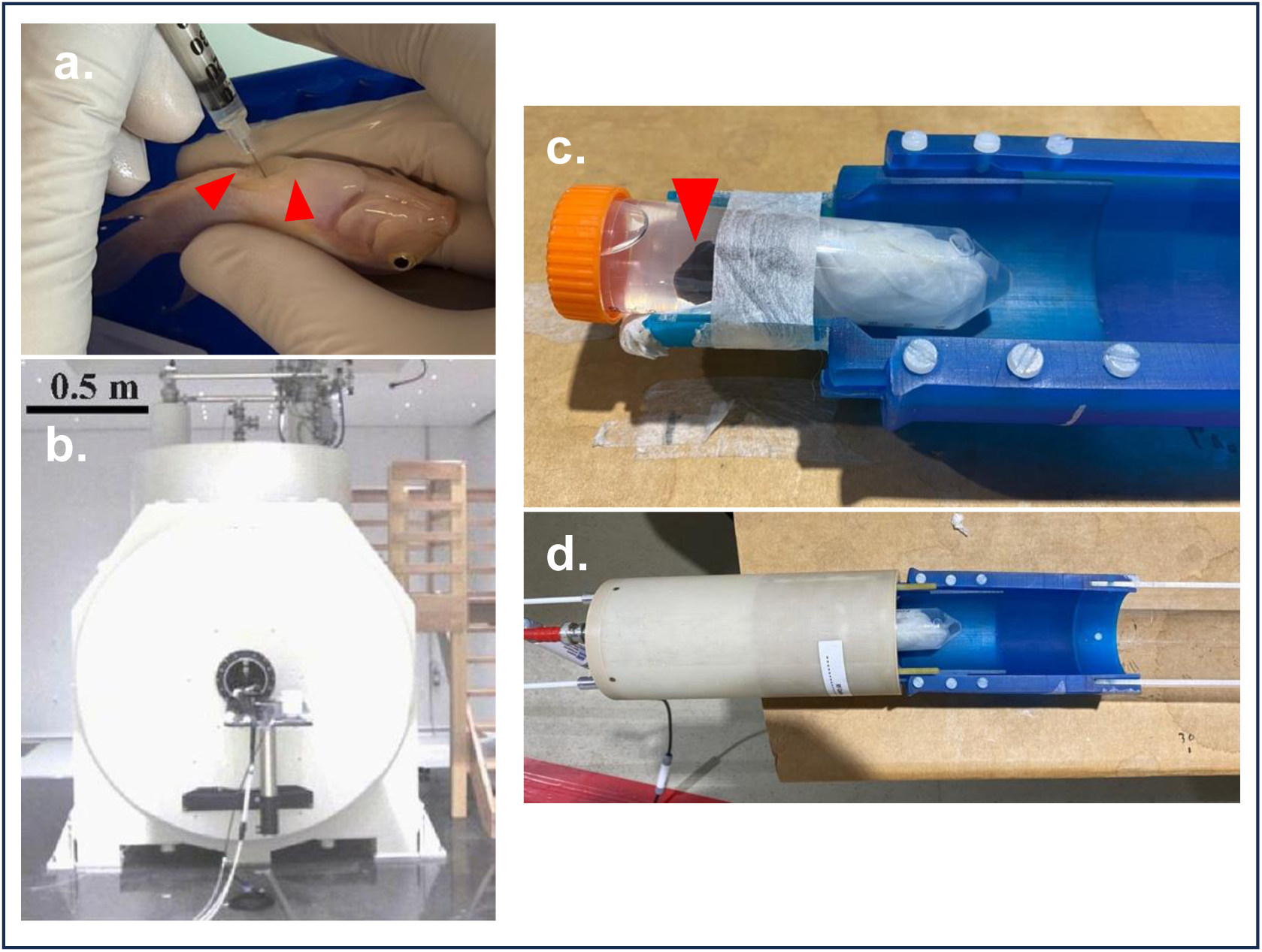
Manganese Enhanced MRI (MEMRI) setup for convict cichlids. a. Intraperitoneal injection of MnCl_2_. The needle is placed between the pelvic fins and the anus, and plunged approximately 3mm into the abdominal wall to reach the intraperitoneal compartment. b. The 17.2 T Bruker Biospin scanner. c. The fish is anesthetized and placed in a 50 mL Falcon tube filled with anesthesia water (MS222, 180 mg/L), its body wrapped in a paper towel to prevent movement. The tube is then taped to the bed of the MRI. The convict cichlid’s head can be seen (red arrowhead). d. The fish is placed inside the RF coil and transferred inside the scanner.

**Figure 2.**
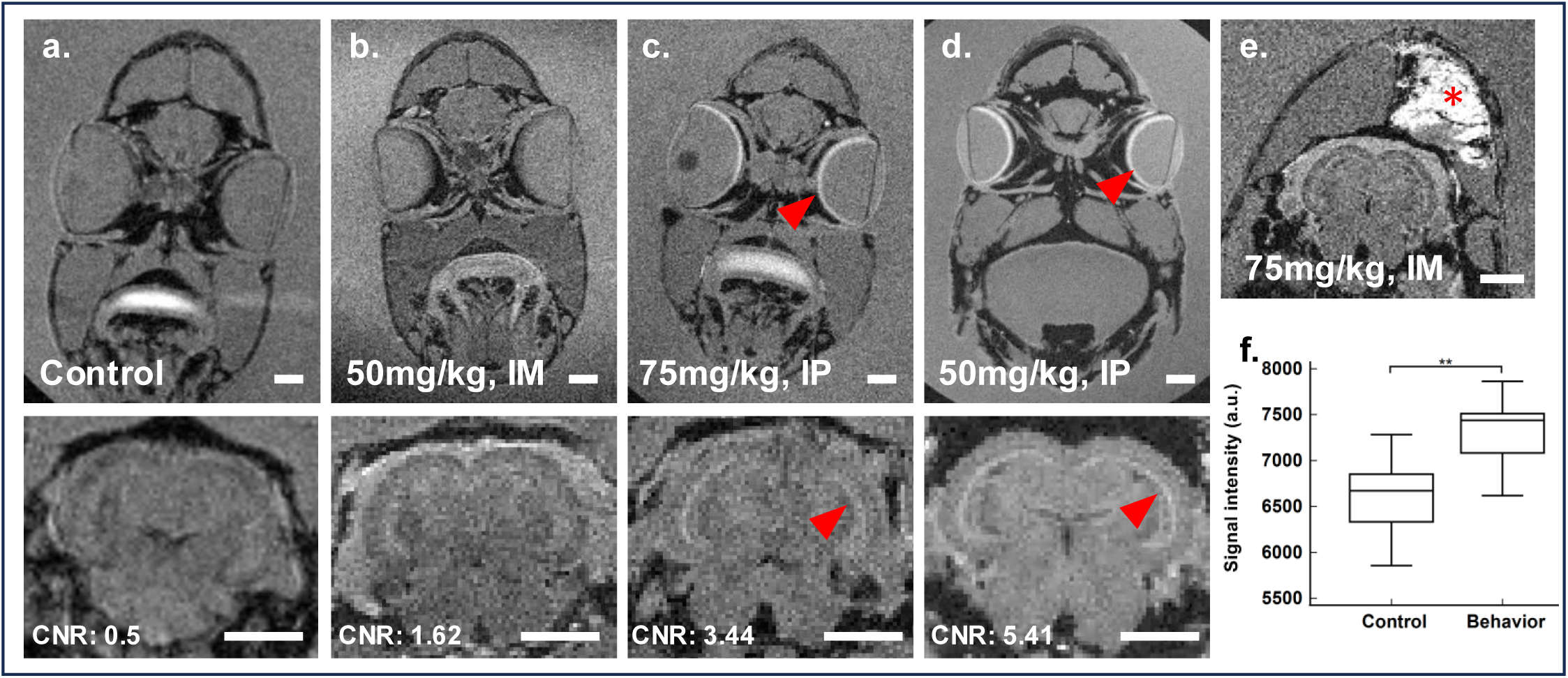
Manganese dosage for MEMRI in convict cichlids. T1 weighted images. a-d: Intraperitoneal injections provide better signal than intramuscular injections for similar manganese concentrations, and 50 mg/kg provides sufficient contrast. Upper pannel: frontal view of the head of convict cichlids at the level of the telencephalon, injected with vehicle (a), 50 mg/kg MnCl_2_ intramuscularly (b), 75 mg/kg MnCl_2_ intraperitonealy (c), and 50 mg/kg MnCl_2_ intraperitonealy (d). No manganese is visible in the retina of (a) and (b), while both (c) and (d) display manganese accumulation in the retina (red arrowhead). Lower pannel: frontal view of the brain at the level of the optic tectum and inferior lobe, showing manganese accumulation in the retinal layer of the optic tectum (red arrowhead) in (c) and (d) Contrast to noise ratio (CNR) is indicated for each image. Acquisition parameters were optimized for (d), which could explain its higher CNR compared to (c). e. Fish injected intramuscularly with 75 mg/kg MnCl_2_. Frontal view at the level of the optic tectum showing a large accumulation of manganese in the left expaxial muscle (red asterisk) where injection was performed, demonstrating the suboptimal kinetics of IM MnCl_2_ injections in ectotherms. f. Mean relative signal intensity in the inferior lobe of the brain of a control group which was only given food (left), and a behavioral group which had to open a box in order to obtain food (right), showing significantly higher manganese accumulation in the behavioral group (p<0.01). Scale bar: 2.5 mm.

Fish were left overnight in their tank to recover from anesthesia and manganese administration. In order to maximize manganese accumulation related to the behavior of interest, as many behavioral sessions as possible should be performed before imaging the animals, and disturbances which might bias the results should be avoided (e.g. unusual noise in the animal house facility). The behavioral task required opening a box to obtain a food reward. We performed two sessions of one hour each the day following the injection, and a final session of one hour right before imaging, two days after the injection. This allowed the animals enough time to recover from the injection, while maximizing manganese accumulation. While manganese clearance is faster in endotherms than in ectotherms^28,35^, manganese accumulation in the brain peaks around 24-48 hours in mice^36^, suggesting that in both endotherms and ectotherms, imaging should be performed within 48 hours of manganese administration.

48 hours following manganese injection, fish were imaged on a 17.2 T imaging system (Bruker Biospin, Fig. 1b). We anesthetized the fish (MS222, 180 mg/L) and placed them in a Falcon 50 mL tube filled with anesthesia water. The animals’ bodies were wrapped in a paper towel to minimize movement within the tube (Fig. 1c), leaving only the head and gills outside the paper towel, and the tube was placed in the coil (Fig. 1d). Using this setup, convict cichlids could remain anesthetized for approximately two hours. Two consecutive T1 weighted images were acquired at an in-plane resolution of 80 µm for a total acquisition time of 52 minutes, after which the fish were ventilated with fresh water to end anesthesia.

Data was processed using the FSL suite of tools^37^. Images were normalized on the retina, which always displayed very high levels of manganese accumulation (Fig. 2). A region of interest (ROI)-based analysis comparing the mean intensity of the brain structure of interest in an experimental group versus a control group can then be used to identify the brain regions involved in the behavior of interest, with significantly higher signal intensity in the behavioral group (Fig. 2f) (Two-way ANOVA and Tukey’s post-hoc test, p<0.01).

Overall, MEMRI enables functional MRI (fMRI) in cichlid fish. Using this technique, it thus becomes possible to non-invasively identify the brain structures involved in any behavior that can be reliably reproduced in the lab.

### High-Intensity Focused Ultrasound (HIFU) protocol guided with MR

To confirm the involvement of a brain structure in a specific behavior as detected with MEMRI, HIFU can be used for lesional studies. Behavioral deficits following the lesion would provide additional evidence of the brain structure’s role in the behavior. In case of non-conventional animal models like cichlids, where a stereotaxic atlas is not available, ultrasound must be combined with MR imaging to accurately target the selected brain region. Therefore, a specific multi-modality protocol was developed (Fig. 3a). This protocol integrates MR sequences to precisely position the ultrasound focal spot and monitor tissue temperature rise, as well as ultrasound sequences to perform thermal ablation.

**Figure 3.**
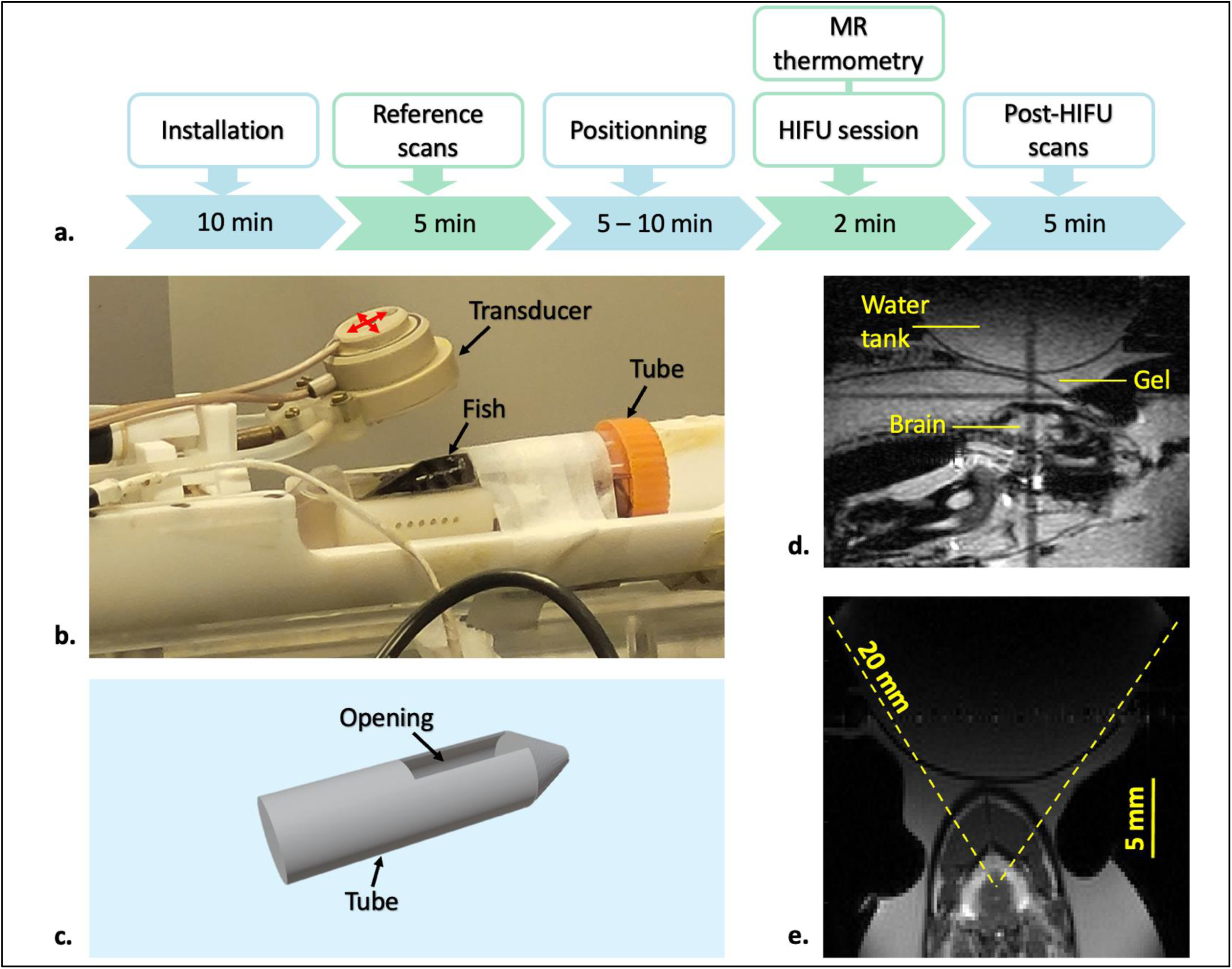
Experimental setup for simultaneous HIFU and MR imaging. a. Workflow of the study with the timeline of each step. b. MR and ultrasound setup, the transducer can move along the two perpendicular directions symbolized by the red arrows. c. The fish was placed in the tube and maintained in water under anesthesia. An opening was created to place the transducer on the fish head. Sagittal T1-weigthed (d.) and coronal T2-weigthed (e.) images.

To enable multi-modal imaging, a dedicated setup was designed to achieve thermal ablation in the targeted region of the fish brain (Fig. 3b). After anesthesia induction, the fish was placed and secured in place with gauze in a Falcon 50 mL tube filed with water with an opening at the top (Fig. 3c). This opening allowed access to the fish’s brain for the ultrasound device while keeping the fish submerged in water. The tube containing the fish was then positioned within a specialized setup that included an RF coil, as well as an ultrasound transducer and its electronics, which were compatible with the MR scanner. The transducer’s position was controlled by motors, enabling movement along two perpendicular directions, as indicated by the red arrows in Fig. 3b^31^.

Two structures were targeted: the cerebellum, as a superficial brain structure, and the inferior lobe, a ventrally located structure. T1- and T2-weighted scans allowed precise positioning of the transducer at the desired location (Fig. 3d). On axial T2-weighted images, two lines were drawn at the MR console (Fig. 3e) to determine the location of the ultrasound focal spot. The transducer was then repositioned, and an additional T2-weighted scan was acquired to confirm that the focal spot was correctly aligned with the cerebellum or inferior lobe.

The MRI-guided HIFU protocol was applied to n=12 convict cichlids. The corpus cerebelli of the cerebellum was selected as the test target for thermal ablation (n=6), as extensive lesions of the corpus cerebelli have been shown to be well tolerated in other teleost fish such as the goldfish^25,38–41^, without impairing swimming and food intake. As our interest is to study cognitive capacity using food-reinforced behaviors, it was important not to destroy “basic” behaviors like swimming and food-intake. Due to its dorsal location and ease of visual examination in dissected brains, the corpus cerebelli was an ideal structure to investigate whether lesions could be induced. Additionally, it provided an opportunity to assess whether the lesions were sufficiently small to target the corpus cerebelli alone, without impacting adjacent regions, thereby preserving the fish’s behavioral integrity.

An MR thermometry acquisition, lasting 1 minute and 55 seconds, was initiated 30 seconds before sonication. The HIFU sequence was then applied for 15 seconds. A temperature increase at the ultrasound focal spot was observed on the thermometry map (Fig. 4a). Tissue temperature over time is shown in Fig. 4b: a sharp temperature increase was detected during ultrasound exposure, peaking at the end of the ultrasound sequence, followed by a gradual decline back to baseline levels.

**Figure 4.**
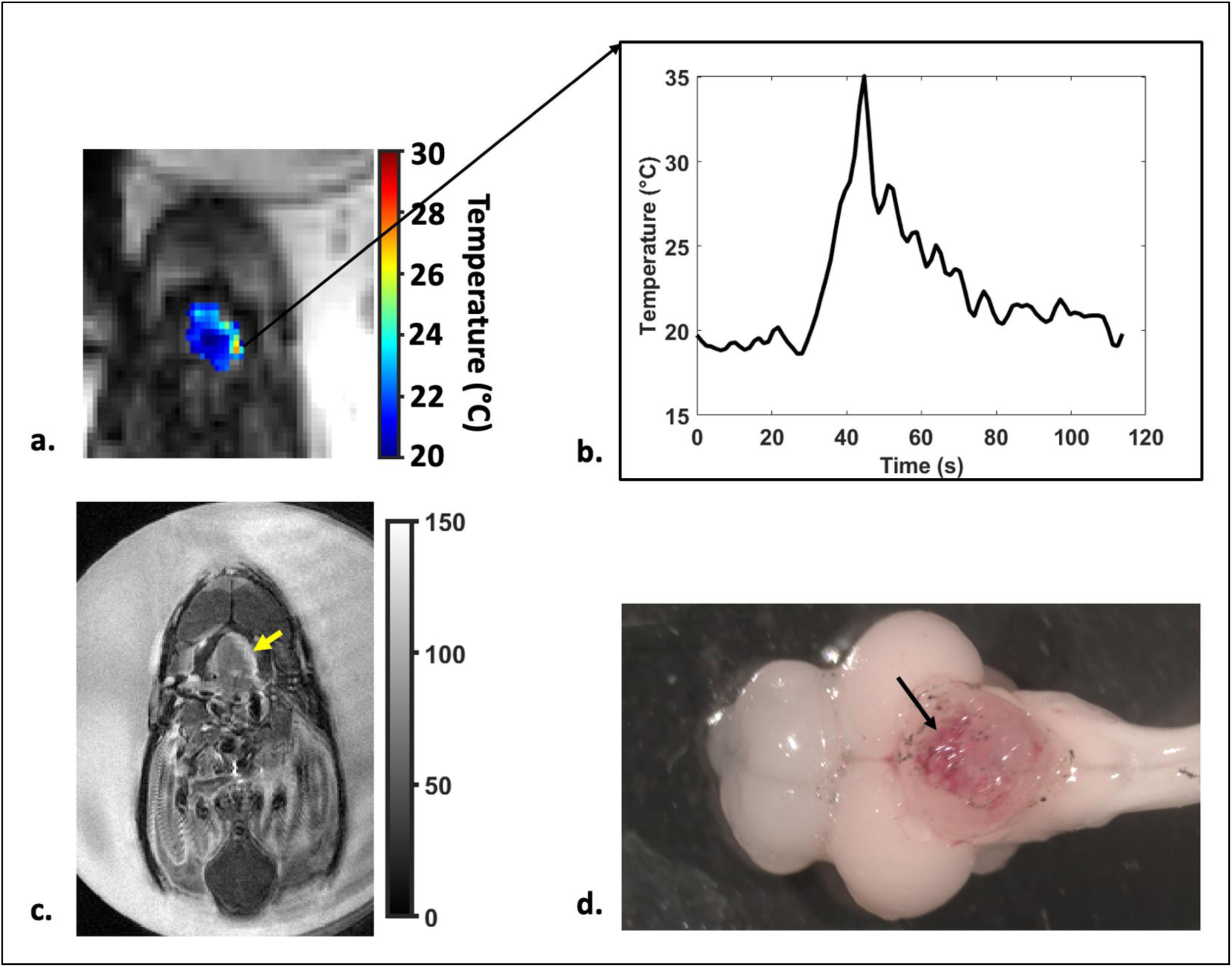
Thermal ablation of the cerebellum. a. Thermometry map of the cerebellum where a temperature rise was detected on the left. b. Temperature evolution over time during MR thermometry scan time. c. An increase of the T2 signal was observed, 48 hours after HIFU protocol, at the location of the temperature rise indicated by the yellow arrow. d. Dorsal view of a lesioned fish brain, showing extensive hemorrhage in the corpus cerebelli (black arrow).

Temperature measurements taken before ultrasound exposure and at the peak of the sequence were compared. An average increase of 18.4 ± 2.2°C was observed between the temperature before and at the end of the sonication (Mann-Whitney test, p<0.05 in all cases).

Thermal ablation induced tissue edema, which was visible on T2-weighted images acquired 48 to 72 hours after the HIFU protocol. In the case presented as an example in Fig. 4c, an increase in the T2 signal, most likely due to edema formation, was observed at the location where a temperature rise had been detected on the thermometry map, corresponding to the ultrasound focal spot (Mann-Whitney test, p<0.05).

This confirms that tissue damage occurred at the intended location (Fig. 4d). Hematoxylin-eosin staining on brain sections was previously used to identify HIFU lesions in rats^32^. However, this method requires somewhat homogeneous cytoarchitecture to distinguish coagulative necrosis and edema from healthy tissue. The small size of teleost brains makes the use of hematoxylin-eosin challenging as changes in healthy cytoarchitecture can be mistaken for lesions and vice-versa. Thus, we visually inspected the brain after dissection following perfusion with 10 % formalin to confirm the presence of a lesion as observed in T2-weighted images. This revealed tissue damage (Fig. 4d) demonstrated by the presence of visible, extensive hemorrhage in the targeted area. There was also a small T2-weighted signal enhancement at the top of the head muscles caused by contact with the transducer setup creating slight heating.

Fish were monitored following HIFU + MR protocol. As expected after a corpus cerebelli lesion, no adverse reactions were observed in any fish 24 hours post-lesion or in the following days (up to a week for n=2 fish, after which they were euthanized) after the experiment. All fish were swimming and feeding normally, indicating that HIFU lesions are compatible with subsequent behavioral experiments.

The fish’s ability to perform a behavioral task requiring goal-directed object manipulation, where the fish had to open a box with their mouth to access a food reward was unaffected. Lesions applied to other brain structures such as the inferior lobe (n=6) resulted in performance deficits in this behavioral task, validating HIFU as a suitable method for lesion studies correlated with behavior in teleost fish.

## DISCUSSION

Overall, MEMRI and HIFU are complementary tools to link behavior and neuroanatomy in non-model species. MEMRI enables MR functional imaging at <100 µm resolution in a non-invasive manner, and manganese is well tolerated in both ecto- and endotherms. A dose of 50 mg/kg seems a good starting point for any vertebrate species and should be sufficient to provide reliable labelling of activated neurons. Higher doses should be tested carefully as they may produce undesirable side-effects on animals and impact their ability to perform their behavioral task. Particular care should be taken to have the animals perform their behavioral task as many times as possible following manganese injection, as it is the accumulation of manganese that is detected in MEMRI. Thus, at least a one-hour session right before imaging is necessary, and more sessions should be performed if possible. Finally, anesthesia must be monitored closely during imaging, as the death of the animal would impact the reliability of the data. Manganese quickly exits cells following death; it is thus crucial to maintain the animals alive for the duration of imaging. For tetrapods, a breathing monitoring system can be used to monitor breathing rate inside the scanner. In fish, we tested progressively longer anesthesia times outside of the scanner to make sure the duration of the imaging was not lethal for the animals, as monitoring breathing during imaging was impossible. We recommend testing anesthesia before starting a MEMRI protocol for species without established anesthesia guidelines. The primary limiting factor of MEMRI is the size of the coil and of the MR scanner bore, which must be large enough to accommodate the animals.

HIFU enables lesion studies without surgery to validate functional imaging findings. HIFU can produce lesions as small as transducer focal spot permits (about 1mm^3^) and is particularly useful for targeting ventrally located structures that might be more difficult to access through surgery. As with MEMRI, anesthesia needs to be carefully managed during HIFU, as the animal must be kept in place when ultrasounds are applied. In our case, besides anesthetizing the fish with MS222, we securely cradled them in squares of gauze to prevent any movement within the tube.

The transducer must be calibrated to determine the delivered acoustic pressure using a hydrophone system^31^. In rodents, high acoustic pressure (between 2.65 and 3.65 MPa) applied for a short duration (under 10 seconds) has been shown to be sufficient to induce necrosis^32^. However, the pressure must be adjusted according to the animal model, as acoustic transmission depends on multiple factors such as skull thickness and body mass, as described in rodents^42^. In this study, the acoustic pressure was adapted to the specific animal model, which has a thin skull but a substantial muscle layer above it. Using a 1.5 MHz central frequency FUS transducer, an acoustic pressure of 5 MPa was sufficient to induce necrosis in the cerebellum. However, for deeper structures such as the inferior lobe, steering is required, and power must be adjusted to maintain the same acoustic pressure range. Steering range, and thus available thermal ablation depth, will depend on the transducer used. Simultaneous thermometry imaging during sonication is essential to monitor heat generation and assess the affected area. Here, the maximum temperature reached was slightly lower than that observed in rodents for necrosis induction. This discrepancy may be due to the orientation of the thermometry slices. Because of animal constraints, slices were aligned parallel to the transducer beam, which can lead to partial volume errors and an underestimation of temperature^43^.

MEMRI is limited in temporal resolution compared to other techniques like BOLD fMRI^44^ and electrophysiology. In addition, in both BOLD and electrophysiology, the animals are their own controls within the same imaging session, with data acquisition starting before the task is initiated, producing baseline data to compare behavioral data to. The problem of BOLD fMRI in animals is that it is difficult to perform behavioral tasks inside the MR scanner: only very few species can be trained to do so. Electrophysiology has the same limitation, and additionally requires the surgical implantation of recording electrodes, which is very difficult in aquatic species as described previously. By contrast to these two techniques, MEMRI can be applied to a much broader ranges of species and behavioral protocols.

Compared to HIFU, optogenetic inactivation of a brain region is reversible and can be switched on and off rapidly, offering high temporal resolution. However, optogenetic inactivation is dependent on gene expression, and often there are no genes which are exclusively expressed in a specific region. Thus, when we want to target a specific region, lesions would be more suitable.

MEMRI and HIFU thus compare favorably to other techniques, especially when dealing with non-conventional model species. In particular, as our pilot study in fish showed, they are extremely well-suited for aquatic species, for which most of the cognitive neuroscience toolkit is unavailable.

In conclusion, MEMRI and HIFU have few limitations regarding the species used, as neither require prior neuroanatomical (e.g., availability of a stereotaxic brain atlas) data to work. This is particularly useful in species with continuously growing nervous systems in adulthood, like fish^45^ or cephalopods^46^, for which stereotaxic brain atlases are not available. Additionally, these techniques are useful for species with large amounts of CSF around the brain, such as reptiles^47^, where traditional stereotaxic surgery may be unreliable. By enabling the study of non-traditional species, MEMRI and HIFU have the potential to open new horizons in our understanding of diverse nervous systems.

## METHODS

### Animals

Sexually mature individuals of both sexes of the convict cichlid *Amatitlania nigrofasciata* measuring between 4 and 7.5 cm in standard length were obtained from a commercial supplier (Aquariofil.com, Nîmes, France). Fish were housed in large 120 to 400L filtered and aerated fresh water communal tanks, with water kept at 25°C. A light/dark cycle of 12/12 hours was maintained throughout the duration of the experiments. A month prior to the start of the MEMRI and HIFU protocols, individuals were isolated for behavioral training in 40L tanks placed next to each other so that visual contact could be maintained between individuals.

48-72 hours (n=10) and up to a week (n=2) after the HIFU protocol, fish were deeply anesthetized (MS222, 300 mg/L) and perfused transcardially with 10% formalin. Following perfusion, the brain was dissected and photographed to visually confirm the location of the lesion.

All procedures were conducted in compliance with the official regulatory standards of the French Government, and the committees (CEA, NeuroPSI) under protocols APAFIS #51934-2024111418122817 v5; APAFIS #55234-2025051210564718 v1 & APAFIS #40422-2023012000331085 v7.

### MnCl_2_ injection, MEMRI preparation and anesthesia

48 hours prior to imaging, fish were anesthetized in MS222 (170 mg/L), weighed, and injected intraperitoneally with a solution of MnCl_2_ in 1M HEPES (pH 7.5, 50 mg/kg) (n=22). Fish were injected with MnCl_2_ under anesthesia (MS222, 170 mg/L) in between the anus and pelvic fins by driving an insulin needle 3 mm into the muscle wall to access the peritoneal space (Fig. 1a). The depth of injection for the size range of individuals used was determined by prior dissection of a fish to measure the thickness of the abdominal wall.

Multiple MnCl_2_ doses were tested, starting with the highest tolerated dose of 75 mg/kg (Fig. 2c). However, some lethality was observed at this dose, as well as at a lower dose of 65 mg/kg. Thus, we settled on a dose of 50 mg/kg. Fish were given 24 hours to recover, then two behavioral sessions were performed 24 hours prior to imaging, spaced approximately 6 hours apart. On the day of imaging, a final behavioral session was performed one hour before imaging. At the end of the session, the animals were anesthetized in MS222 (180 mg/L), then transferred in a 50 mL Falcon tube filed with fresh anesthesia water. Depending on the size of the animal, a paper towel was wrapped around the fish to keep it fixed inside the tube.

At the end of acquisition, fish were ventilated with fresh water until voluntary movement started again, then transferred back into their housing tanks. The procedure from beginning of anesthesia to end of acquisition lasted approximately two hours. No toxicity from the manganese injection was observed, up to a year after injection.

### HIFU parameters

An MR-compatible, 8-element focused ultrasound transducer with a central frequency of 1.5 MHz, a geometrical depth of 20 mm and a diameter of 30 mm (Imasonic, Besançon, France) was used. The transducer was coupled to the animals via a balloon filled with degassed water, sealed with a latex membrane. Water was degassed for 30 minutes prior to each experiment. The transducer was mounted on a mobile stage, and its position could be adjusted from outside the magnet using MR-compatible motors^31^. Motor movements and ultrasound parameters were controlled by dedicated software (Thermoguide®, Image Guided Therapy, Pessac, France). Fish were sonicated through an intact scalp using 800 ms pulses delivered once per second over a 15-second period, with an estimated focal acoustic pressure of 6 MPa in the brain. Acoustic pressure values were determined from prior transducer calibration in a degassed water tank using a hydrophone (HGL-0200, preamplifier AG-2010, Onda Corporation, USA). HIFU experiments were conducted on 12 fish, with MR guidance used to target the cerebellum (n=6) or the inferior lobe (n=6).

### MR acquisitions

MEMRI acquisitions were performed on a 17.2 T Bruker imaging system (Bruker Biospin) equipped with an Avance III console running Paravision 6.0.1 using a 45 mm diameter birdcage transmit/receive volume coil (Rapid Biomedical) on n=22 fish. Two consecutive T1 weighted images were acquired with the following parameters: echo time (TE)/repetition time (TR) = 2.8/250ms, flip angle = 90°, 24 slices, slice thickness= 0.2 mm, in plane resolution = 0.08 x 0.08 mm^2^, acquisition time = 26 min.

HIFU experiments were conducted under MR guidance using a 7 T/90 mm Pharmascan (Bruker Biospin) with a dedicated ultrasound single-loop radiofrequency coil^31^, whose diameter was wide enough for the ultrasound beam to pass through it and for extensive displacement of the transducer above the fish head. First, anatomical T2-weighted MR images were acquired by using a rapid acquisition with relaxation enhancement spin echo sequence (RARE). The acquisition parameters included effective TE/TR=20.75/3000ms, RARE factor = 8, in plane resolution = 0.25 x 0.25 mm^2^, filed of view = 32 x 32 mm^2^, 19 axial slices, slice thickness = 1 mm, total scan time = 3 min 12 s. When positioning slices, care was taken to include the transducer border in the field of view, as T2-weighted images were used to accurately position the ultrasound spot to the target area. The T2-weighted acquisition was repeated with the same parameters after thermal ablation.

To monitor the temperature during thermal ablation an MR thermometry acquisition was performed. A standard Fast Low Angle Shot (FLASH) sequence was used with TE/TR= 3.5/10 ms, and a flip angle of 30°. An image was generated every 1.28 s with a resolution of 0.5 x 0.5 x 3 mm^3^, for a field of view of 32 x 32 mm^2^. One single slice was acquired in axial orientation for a scan time of 1 min 55 s. The MR thermometry sequence was started 30 s before the ultrasound exposure. All experiments were conducted in a room with a temperature of 20°C.

A high-resolution T2-weighted RARE acquisition was performed 48 to 72 hours after thermal ablation. The following parameters were used: TE/TR = 8/1500 ms, field of view = 28 x 28 mm^2^, spatial resolution = 0.2 x 0.2 x 0.2 mm^3^, 15 slices acquired in axial orientation.

### MR data processing

MEMRI data was processed using the FSL suite of tools^37^. The two T1 sequences were averaged to produce a single, higher signal to noise ratio image. To normalize the signal intensity across individuals, the mean intensity of the retina of each individual was measured, as the retina was reliably labeled with manganese in all fish. The MRI data was divided by this value and multiplied by an arbitrary value of 10,000 to obtain normalized images. Contrast to noise ratio (CNR) was calculated by dividing the difference of the mean intensity in the retina with the rest of the eye by the standard deviation of the water surrounding the fish. FSLeyes was used to draw ROIs and measure mean intensity for each ROI.

For HIFU MR thermometry, images acquired before ultrasound exposure were used as baseline. After phase unwrapping, the phase image acquired before ultrasound exposure (baseline) was subtracted from the unwrapped phase images acquired during and after ultrasound exposure. This phase difference was then used to generate temperature maps. The temperature rise (ΔT, in degrees Celsius) was calculated using the following equation

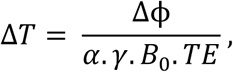

Where Δϕ is the phase difference in radians, 𝑇𝐸 is the echo time, 𝐵_!_ is the strength of the magnetic field, 𝛾 is the gyromagnetic ratio and 𝛼 is the temperature sensitivity coefficient at −0.01ppm/°C. For B_0_ = 7 Tesla, a value of 3.1Hz/°C was used for 𝛼 ^48^. Using T2-weighted images obtained 48 to 72 hours after ultrasound exposure, an ROI was placed in the approximate area where a temperature increase had been detected in MR thermometry. A second ROI was placed in a brain region not exposed to ultrasound. The signal to noise ratio was then estimated in both ROIs.

### Statistics

Comparison between the MEMRI experimental and control group in Fig. 2f was performed using a two-way ANOVA and post-hoc Tukey test in R Studio v.2023.12.1+402. Comparison between temperatures measured before and during ultrasound exposure were performed with the Mann-Whitney test. Additionally, a comparison of the signal to noise ratio estimated between the brain areas exposed and non-exposed to ultrasound was performed. Statistical significance was considered for P values <0.05. Statistical analysis was performed using Medcalc version 20.014 (Medcalc, Ostend, Belgium).

## DATA AVAILABILITY

Data reported in this paper will be shared by the corresponding authors upon request. This paper does not report original code.

## ACKNOWLEDGEMENTS

We thank Anthony Novell, Corentin Cornu and the BioMaps laboratory (Paris-Saclay University, CEA, CNRS, Inserm, BioMaps, Service Hospitalier Frédéric Joliot, Orsay, France) for their help in calibrating the ultrasound transducer.

This study was supported by INSB/CNRS Call “Diversity of Biological Mechanisms” and ANR EVONECTOME (ANR-23-CE37-0021).

## AUTHOR CONTRIBUTIONS

Conceptualization, K.Y. and P.E.; methodology, L.C., G.P., P.E., B.L., K.Y.; funding acquisition and supervision, K.Y. and L.C.; validation and visualization, K.Y., P.E., L.C., G.P.; first draft of manuscript, P.E. and G.P.; all authors contributed to data analysis, interpretation and revision of the manuscript.

## MATERIALS & CORRESPONDENCE

Correspondence and requests should be addressed to Luisa Ciobanu or Pierre Estienne.

